# The pathogenicity of *PSEN2* variants is tied to Aβ production and homology to *PSEN1*

**DOI:** 10.1101/2024.06.22.600217

**Authors:** Lei Liu, Stephanie A. Schultz, Adriana Saba, Hyun-Sik Yang, Amy Li, Dennis J. Selkoe, Jasmeer P. Chhatwal

## Abstract

**INTRODUCTION:** Though recognized as a potential cause of Autosomal Dominant Alzheimer’s Disease, the pathogenicity of many *PSEN2* variants remains uncertain. We compared Aβ production across all missense *PSEN2* variants in the Alzforum database and, when possible, to corresponding *PSEN1* variants.

**METHODS:** We expressed 74 *PSEN2* variants, 21 of which had homologous *PSEN1* variants with the same amino acid substitution, in HEK293 cells lacking PSN1/2. Aβ production was compared to age at symptom onset (AAO) and between homologous *PSEN1/2* variants.

**RESULTS:** Aβ42/40 and Aβ37/42 ratios were associated with AAO across *PSEN2* variants, strongly driven by *PSEN2* variants with *PSEN1* homologs. *PSEN2* AAO was 18.3 years later compared to *PSEN1* homologs. Aβ ratios from *PSEN1*/*2* homologs were highly correlated, suggesting a similar mechanism of γ-secretase dysfunction.

**DISCUSSION:** The existence of a *PSEN1* homolog and patterns of Aβ production are important considerations in assessing the pathogenicity of previously-reported and new *PSEN2* variants.

## 1. INTRODUCTION

Nearly 500 pathogenic variants leading to Autosomal Dominant Alzheimer’s disease (ADAD) have been identified in *PSEN1*, *PSEN2*, and *APP* (*1–3*). Though all these variants are considered highly penetrant, the age of symptom onset (AAO) within families and between pathogenic variants varies widely, encompassing a span of 50+ years (*4, 5*). The largest number of pathogenic variants described thus far are in *PSEN1* and constitute the most common cause of ADAD, accounting for 70% to 80% of cases (*1*). Presenilin 1 (PSN1), encoded by *PSEN1*, forms the catalytic core of the γ-secretase complex, an enzyme complex that directly mediates the production of longer, aggregation-prone Aβ peptides relative to shorter, non-aggregating peptides (*6–14*). A smaller number of ADAD-causing pathogenic variants have been identified in *PSEN2* (*2*), which has many similarities to *PSEN1* and its encoded protein, presenilin 2 (PSN2), can similarly serve as the catalytic core of a functional γ-secretase complex (*15–17*). Despite many similarities, PSN2 and PSN1 are thought to have subtle variations in their sub-cellular localization (*18, 19*) and overall expression levels (*20–23*). Intriguingly, the average AAO in individuals bearing *PSEN2* variants is substantially older than that seen in carriers of *PSEN1* variants (*24, 25*), though reasons for this striking dissimilarity are not well understood. Recent studies have compared the Aβ production profiles between wild-type PSN1 and PSN2 (*26*), but there have been no previous studies comparing conserved mutations across *PSEN1* and *PSEN2*.

Recent work from our group and others has shown that the clinical and biomarker changes seen across the ADAD disease course in *PSEN1* variant carriers may be due in large part to the extent to which individual variants impact the production of aggregation-prone Aβ species (Aβ42 and 43) as compared to shorter, less-aggregation prone species (Aβ37 and 38) (*24, 27–32*). Indeed, quantitative analysis of amyloid precursor protein (APP) processing by individual *PSEN1* variants reveals strong associations between the ratio of Aβ42/Aβ40 and/or Aβ37/42 production with the AAO and rate of cognitive decline across more than 160 examined *PSEN1* variants (*30*). Beyond AAO, we have recently shown strong associations between the ratio of short-to-long Aβ species 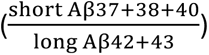 and a wide set of *in vivo* imaging and biofluid markers of neurodegeneration, tau, and β-amyloid pathology in *PSEN1* pathogenic variant carriers (*27, 33*). Broadly, the findings across *PSEN1* variants strongly suggest that variant-level differences in γ-secretase dysfunction are critical determinants of the clinical and biomarker course of ADAD.

The associations between γ-secretase enzymatic abnormality, Aβ production, and pathogenicity of ADAD has not been fully established as regards *PSEN2* variants. In addition to informing clinical care for people with suspected ADAD-causing variants and the conduct of ongoing clinical trials in ADAD, understanding the link between altered γ-secretase function and the pathogenicity of *PSEN2* variants is particularly critical as there are many variants reported in *PSEN2* that are reported as questionably pathogenic, unclassified, or of uncertain significance. In addition to AD, variants in *PSEN2* have also been linked with other neurological disorders not classically associated with mutations in the *PSEN1 (*as reviewed by Cai and colleagues (*16*)), such as dementia with Lewy bodies (*34*), frontotemporal dementia (*35*), and Parkinson’s disease dementia (*36, 37*). In this context, the systematic evaluation of *PSEN2* variants may help to refute or support the pathological significance of individual variants in clinical and research settings, particularly when limited family history is available. In this context, we provide a comprehensive examination of γ-secretase function and Aβ production across all non-synonymous, missense *PSEN2* variants currently listed in the Alzforum database, elucidating the differential pathogenicity of similar variants in *PSEN1* as compared to *PSEN2* and to inform clinical evaluation for people bearing *PSEN2* variants of uncertain pathogenicity.

## 2. METHODS

### 2.1 Generation of human PSEN1 and PSEN2 expression vectors

The coding sequence of the wild type human *PSEN1* and *PSEN2* were subcloned into the pcDNA3.1 vector, which was then used to generate vectors expressing specific mutations. Polymerase chain reaction (PCR) amplified the template into two DNA fragments with an overlapping sequence containing the target locus to introduce mutations. Overlap PCR was done to generate the whole open-reading frame containing the mutations. PCR products were subcloned into the same pcDNA3.1 vector for expression under a human cytomegalovirus promoter. Vectors were sequenced from 5’ and 3’ ends to confirm successful mutagenesis.

Using the Alzforum database (*2*), we identified all *PSEN2* missense variants reported (a total of 74 at the start of this study in September 2023), including all of those listed as pathogenic, benign, not classified, associated with FTD or Parkinson’s disease dementia, or of uncertain significance (VUS; see **Table 1** for a list of variants). Due to the model system used, *PSEN2* variants in which the wild-type amino acid was not altered (i.e., synonymous mutations) or where a frameshift mutation was present were not included in the present study. We identified 21 *PSEN2* variants for which the same amino acid substitution was present in a previously-reported, pathogenic variant in *PSEN1*. For the purposes of discussion, we refer to similar substitutions at consonant residues of PSN1 and PSN2 as homologous mutations in *PSEN1* and *PSEN2*. Conversely, *PSEN2* variants without a known *PSEN1* counterpart will be referred to as non-homologous.

**TABLE 1.**
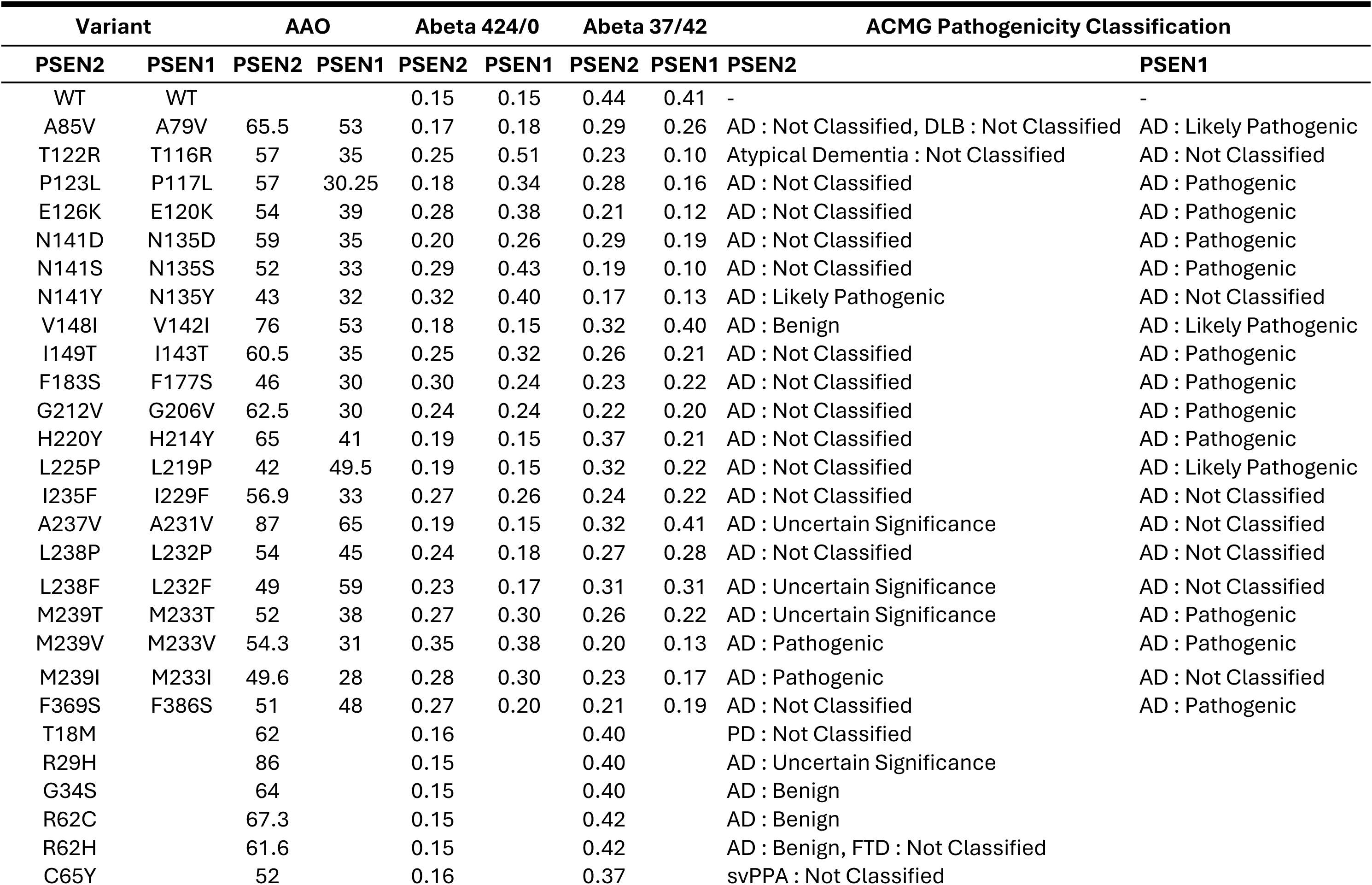
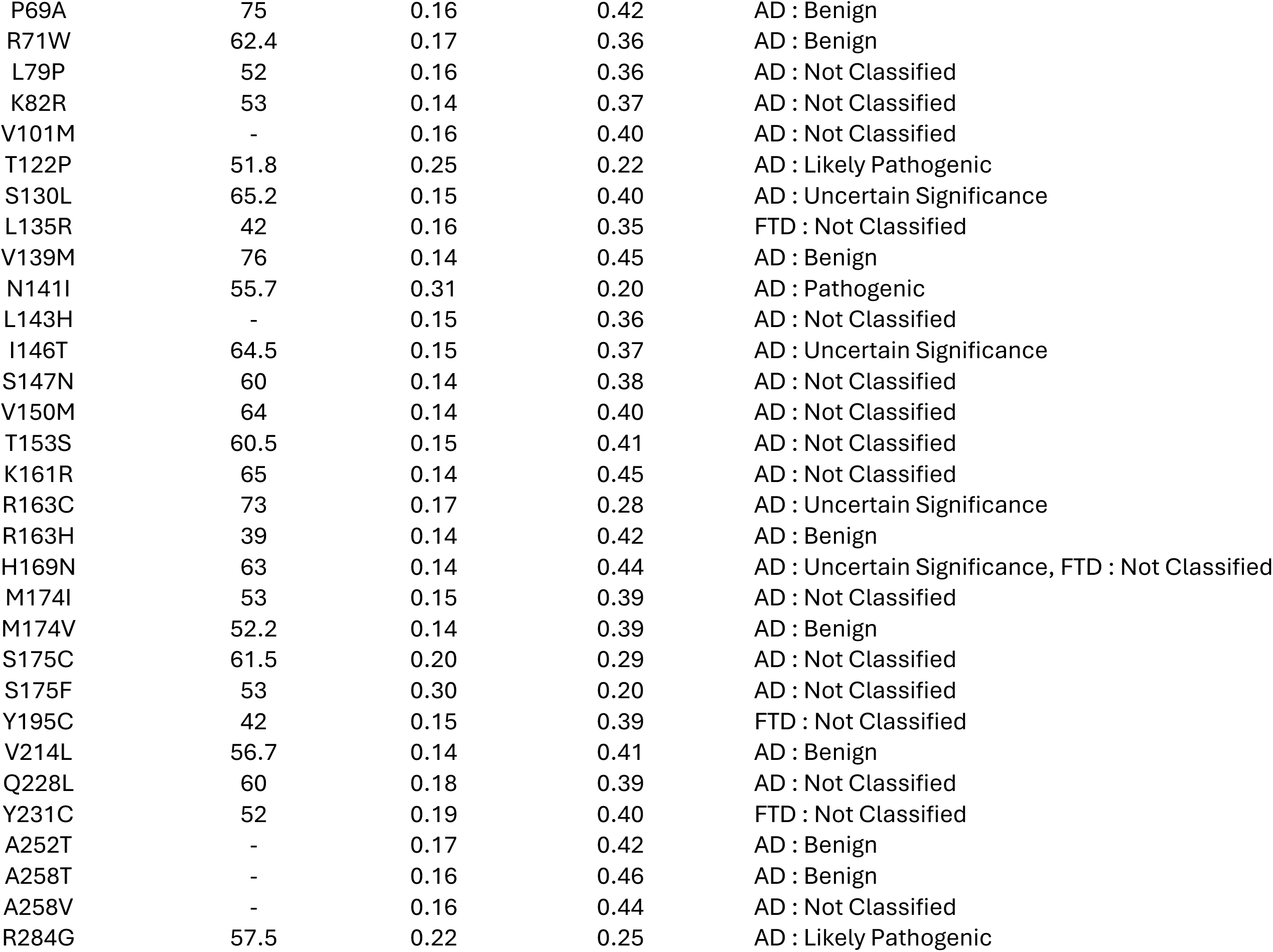
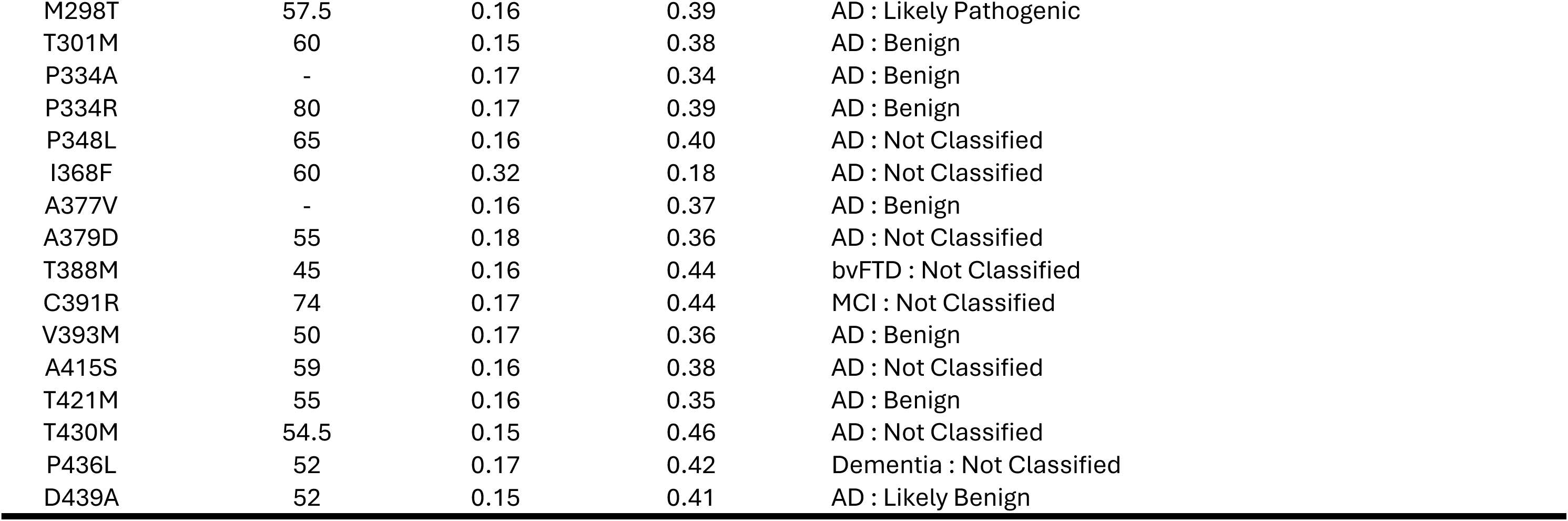
Description of *PSEN1* and *PSEN2* variants characterized.

| Variant |  | AAO |  | Abeta 424/0 |  | Abeta 37/42 |  | ACMG Pathogenicity Classification |  |
| --- | --- | --- | --- | --- | --- | --- | --- | --- | --- |
| PSEN2 | PSEN1 | PSEN2 | PSEN1 | PSEN2 | PSEN1 | PSEN2 | PSEN1 | PSEN2 | PSEN1 |
| WT | WT |  |  | 0.15 | 0.15 | 0.44 | 0.41 | - | - |
| A85V | A79V | 65.5 | 53 | 0.17 | 0.18 | 0.29 | 0.26 | AD : Not Classified, DLB : Not Classified | AD : Likely Pathogenic |
| T122R | T116R | 57 | 35 | 0.25 | 0.51 | 0.23 | 0.10 | Atypical Dementia : Not Classified | AD : Not Classified |
| P123L | P117L | 57 | 30.25 | 0.18 | 0.34 | 0.28 | 0.16 | AD : Not Classified | AD : Pathogenic |
| E126K | E120K | 54 | 39 | 0.28 | 0.38 | 0.21 | 0.12 | AD : Not Classified | AD : Pathogenic |
| N141D | N135D | 59 | 35 | 0.20 | 0.26 | 0.29 | 0.19 | AD : Not Classified | AD : Pathogenic |
| N141S | N135S | 52 | 33 | 0.29 | 0.43 | 0.19 | 0.10 | AD : Not Classified | AD : Pathogenic |
| N141Y | N135Y | 43 | 32 | 0.32 | 0.40 | 0.17 | 0.13 | AD : Likely Pathogenic | AD : Not Classified |
| V148I | V142I | 76 | 53 | 0.18 | 0.15 | 0.32 | 0.40 | AD : Benign | AD : Likely Pathogenic |
| I149T | I143T | 60.5 | 35 | 0.25 | 0.32 | 0.26 | 0.21 | AD : Not Classified | AD : Pathogenic |
| F183S | F177S | 46 | 30 | 0.30 | 0.24 | 0.23 | 0.22 | AD : Not Classified | AD : Pathogenic |
| G212V | G206V | 62.5 | 30 | 0.24 | 0.24 | 0.22 | 0.20 | AD : Not Classified | AD : Pathogenic |
| H220Y | H214Y | 65 | 41 | 0.19 | 0.15 | 0.37 | 0.21 | AD : Not Classified | AD : Pathogenic |
| L225P | L219P | 42 | 49.5 | 0.19 | 0.15 | 0.32 | 0.22 | AD : Not Classified | AD : Likely Pathogenic |
| I235F | I229F | 56.9 | 33 | 0.27 | 0.26 | 0.24 | 0.22 | AD : Not Classified | AD : Not Classified |
| A237V | A231V | 87 | 65 | 0.19 | 0.15 | 0.32 | 0.41 | AD : Uncertain Significance | AD : Not Classified |
| L238P | L232P | 54 | 45 | 0.24 | 0.18 | 0.27 | 0.28 | AD : Not Classified | AD : Not Classified |
| L238F | L232F | 49 | 59 | 0.23 | 0.17 | 0.31 | 0.31 | AD : Uncertain Significance | AD : Not Classified |
| M239T | M233T | 52 | 38 | 0.27 | 0.30 | 0.26 | 0.22 | AD : Uncertain Significance | AD : Pathogenic |
| M239V | M233V | 54.3 | 31 | 0.35 | 0.38 | 0.20 | 0.13 | AD : Pathogenic | AD : Pathogenic |
| M239I | M233I | 49.6 | 28 | 0.28 | 0.30 | 0.23 | 0.17 | AD : Pathogenic | AD : Not Classified |
| F369S | F386S | 51 | 48 | 0.27 | 0.20 | 0.21 | 0.19 | AD : Not Classified | AD : Pathogenic |
| T18M |  | 62 |  | 0.16 |  | 0.40 |  | PD : Not Classified |  |
| R29H |  | 86 |  | 0.15 |  | 0.40 |  | AD : Uncertain Significance |  |
| G34S |  | 64 |  | 0.15 |  | 0.40 |  | AD : Benign |  |
| R62C |  | 67.3 |  | 0.15 |  | 0.42 |  | AD : Benign |  |
| R62H |  | 61.6 |  | 0.15 |  | 0.42 |  | AD : Benign, FTD : Not Classified |  |
| C65Y |  | 52 |  | 0.16 |  | 0.37 |  | svPPA : Not Classified |  |

|  |  |  |  |  |
| --- | --- | --- | --- | --- |
| P69A | 75 | 0.16 | 0.42 | AD : Benign |
| R71W | 62.4 | 0.17 | 0.36 | AD : Benign |
| L79P | 52 | 0.16 | 0.36 | AD : Not Classified |
| K82R | 53 | 0.14 | 0.37 | AD : Not Classified |
| V101M | - | 0.16 | 0.40 | AD : Not Classified |
| T122P | 51.8 | 0.25 | 0.22 | AD : Likely Pathogenic |
| S130L | 65.2 | 0.15 | 0.40 | AD : Uncertain Significance |
| L135R | 42 | 0.16 | 0.35 | FTD : Not Classified |
| V139M | 76 | 0.14 | 0.45 | AD : Benign |
| N141I | 55.7 | 0.31 | 0.20 | AD : Pathogenic |
| L143H | - | 0.15 | 0.36 | AD : Not Classified |
| I146T | 64.5 | 0.15 | 0.37 | AD : Uncertain Significance |
| S147N | 60 | 0.14 | 0.38 | AD : Not Classified |
| V150M | 64 | 0.14 | 0.40 | AD : Not Classified |
| T153S | 60.5 | 0.15 | 0.41 | AD : Not Classified |
| K161R | 65 | 0.14 | 0.45 | AD : Not Classified |
| R163C | 73 | 0.17 | 0.28 | AD : Uncertain Significance |
| R163H | 39 | 0.14 | 0.42 | AD : Benign |
| H169N | 63 | 0.14 | 0.44 | AD : Uncertain Significance, FTD : Not Classified |
| M174I | 53 | 0.15 | 0.39 | AD : Not Classified |
| M174V | 52.2 | 0.14 | 0.39 | AD : Benign |
| S175C | 61.5 | 0.20 | 0.29 | AD : Not Classified |
| S175F | 53 | 0.30 | 0.20 | AD : Not Classified |
| Y195C | 42 | 0.15 | 0.39 | FTD : Not Classified |
| V214L | 56.7 | 0.14 | 0.41 | AD : Benign |
| Q228L | 60 | 0.18 | 0.39 | AD : Not Classified |
| Y231C | 52 | 0.19 | 0.40 | FTD : Not Classified |
| A252T | - | 0.17 | 0.42 | AD : Benign |
| A258T | - | 0.16 | 0.46 | AD : Benign |
| A258V | - | 0.16 | 0.44 | AD : Not Classified |
| R284G | 57.5 | 0.22 | 0.25 | AD : Likely Pathogenic |
| M298T | 57.5 | 0.16 | 0.39 | AD : Likely Pathogenic |
| T301M | 60 | 0.15 | 0.38 | AD : Benign |
| P334A | - | 0.17 | 0.34 | AD : Benign |
| P334R | 80 | 0.17 | 0.39 | AD : Benign |
| P348L | 65 | 0.16 | 0.40 | AD : Not Classified |
| I368F | 60 | 0.32 | 0.18 | AD : Not Classified |
| A377V | - | 0.16 | 0.37 | AD : Benign |
| A379D | 55 | 0.18 | 0.36 | AD : Not Classified |
| T388M | 45 | 0.16 | 0.44 | bvFTD : Not Classified |
| C391R | 74 | 0.17 | 0.44 | MCI : Not Classified |
| V393M | 50 | 0.17 | 0.36 | AD : Benign |
| A415S | 59 | 0.16 | 0.38 | AD : Not Classified |
| T421M | 55 | 0.16 | 0.35 | AD : Benign |
| T430M | 54.5 | 0.15 | 0.46 | AD : Not Classified |
| P436L | 52 | 0.17 | 0.42 | Dementia : Not Classified |
| D439A | 52 | 0.15 | 0.41 | AD : Likely Benign |

### 2.2 Pathogenicity classification and AAO of variants

Pathogenicity classification of *PSEN2* and *PSEN1* variants were determined using ACMG-AMP guidelines and further sub-classified into 3 groups: Pathogenic AD (AD: Pathogenic or AD: likely pathogenic classification), Benign/VUS AD, and Not Classified. AAO was determined from literature search and was available for 67 of 74 *PSEN2* variants and 21 of 21 of *PSEN1* variants.

### 2.3 Tissue culture and transfection of adherent cells

Adherent human embryonic kidney 293 (HEK) cells were cultured in complete growth medium: Dulbecco’s Modified Eagle’s Medium (DMEM) supplemented with 10% fetal bovine serum (FBS), 10 units/ml penicillin, and 10 µg/ml streptomycin. Adherent HEK cells were seeded in a 48-well plate at a density of 200,000 cells per well for transfection. Transfection was carried out with PEI Max. Cells were incubated for 48 hours before medium were harvested for ELISA and western blots.

### 2.4 Generation of *PSEN1/2* dKO cell line with stable human APP expression

The HEK293 *PSEN1/2* dKO cells (*30*) were transduced with a lentivirus expressing human APP-695 with a C-terminal GFP tag. A single clone was isolated with serial dilution. Expression of APP-695-GFP was confirmed with western blot and ELISA.

### 2.5 Aβ immunoassay (MSD)

Conditioned media from cultured cells was diluted with 1% BSA in the wash buffer (TBS supplemented with 0.05% Tween). For Aβ 37/40/42 assays, each well of an uncoated 96-well multi-array plate (MesoScale Discovery, #L15XA-3) was coated with 30 μL of a PBS solution containing 3 μg/mL of m266 capture antibody to the mid-region of soluble Aβ (original gift of P. Seubert, Elan, plc) and incubated at room temperature overnight. We prepared detection antibody solutions containing biotinylated monoclonal antibodies specifically recognizing the C-terminal residue of each Aβ isoform, as well as 100 ng/mL Streptavidin Sulfo-TAG (MesoScale Discovery, #R32AD-5), and 1% BSA diluted in the wash buffer. Following overnight incubation with the capture antibody, 25 μL/well of the sample, followed by 25 μL/ well of detection antibody solution was incubated for 2 hours at RT with shaking at > 300 rpm, with washing of wells with wash buffer (TBS supplemented with 0.05% Tween) between incubations. The plate was read and analyzed according to the MSD manufacturer’s manual. Aβ peptide ratios of 37/42 and 42/40 were chosen *a priori* for analyses.

### 2.6 Statistical analysis

All statistical analyses were performed using R studio (Version 2021.09.0). Pearson correlations were performed to examine (1) the association between AAO and Aβ42/40 or Aβ37/42 in PSEN2 variants, within the full *PSEN2* sample with AAO available (N = 67) and separately in *PSEN2* homologous variants (N = 21) and *PSEN2* non-homologous variants (N = 46); (2) the relationship between AAO of *PSEN1* Homolog variants and AAO of *PSEN2* Homolog variants (N = 21); and (3) the relationship between Aβ42/40 or Aβ37/42 levels in *PSEN1* and *PSEN2* Homolog variants (N = 21). T-tests were performed to examine 1) the difference in AAO between *PSEN1* (N = 21) and *PSEN2* variants (N = 74), 2) within *PSEN2* variants, the difference in AAO among variants classified as AD: pathogenic or AD: likely pathogenic compared (N = 7) to variants with other pathogenicity classifications (AD: benign/AD: VUS and not characterized; N = 60). A linear regression model with the interaction of AAO and *PSEN1/2* grouping interaction as the term of interest was performed to examine how the association between AAO and cell-based Aβ42/40 or Aβ37/42 levels differ between homologous variants in *PSEN1* vs *PSEN2*.

## 3. RESULTS

### 3.1 Pathogenicity classifications of *PSEN2* variants

There was a wide range of ACMG-AMP classifications across the 74 *PSEN2* variants (**Table 1 and Supplemental Table 1**). All homologous *PSEN1* variants characterized were classified as either AD: pathogenic/ AD: likely pathogenic (67%) or AD: Not classified (33%). Though the large majority of *PSEN2* variants were not definitively classified, we observed that *PSEN2* variants with an identified homologous *PSEN1* variant were more likely to be classified as pathogenic (i.e., listed as “AD: pathogenic” or “AD: likely pathogenic” variants in the Alzforum database) compared to those with no known *PSEN1* homologous variant (14% and 8%; **Supplemental Figure 1**). Additionally, *PSEN2* variants classified as AD pathogenic had earlier AAO compared to other variants (t (12) = 2.68 and p = 0.020).

### 3.2 Aβ production patterns are associated with AAO in PSEN2 variants

We observed an association between AAO and the cell-based production ratios Aβ42/40 (r (65) = −0.272 and p = 0.026; **Figure 2A**) and Aβ37/42 (r (65) = 0.251 and p = 0.040; **Figure 2B**) across *PSEN2* variants. However, in examining these data more closely, the associations between Aβ ratios and AAO were completely driven by *PSEN2* variants with a known *PSEN1* homolog (“homologous variants”; Aβ42/40: r (19) = −0.530 and p 0.014; Aβ37/42: r (19) = 0.490 and p = 0.024; **Figure 2C,D**). No such relationship was observed in non-homologous *PSEN2* variants (Aβ42/40: r (44) = −0.105 and p = 0.487; Aβ37/42: r (44) = 0.147 and p = 0.328; **Figure 2E,F**). Further, we observed that a majority of non-homologous *PSEN2* variants had Aβ42/40 and Aβ37/42 ratios that were similar to that observed from wild-type *PSEN2 (*e.g., Aβ42/40 close to 0.1), arguing against the pathogenicity of these variants.

**FIGURE 1.**
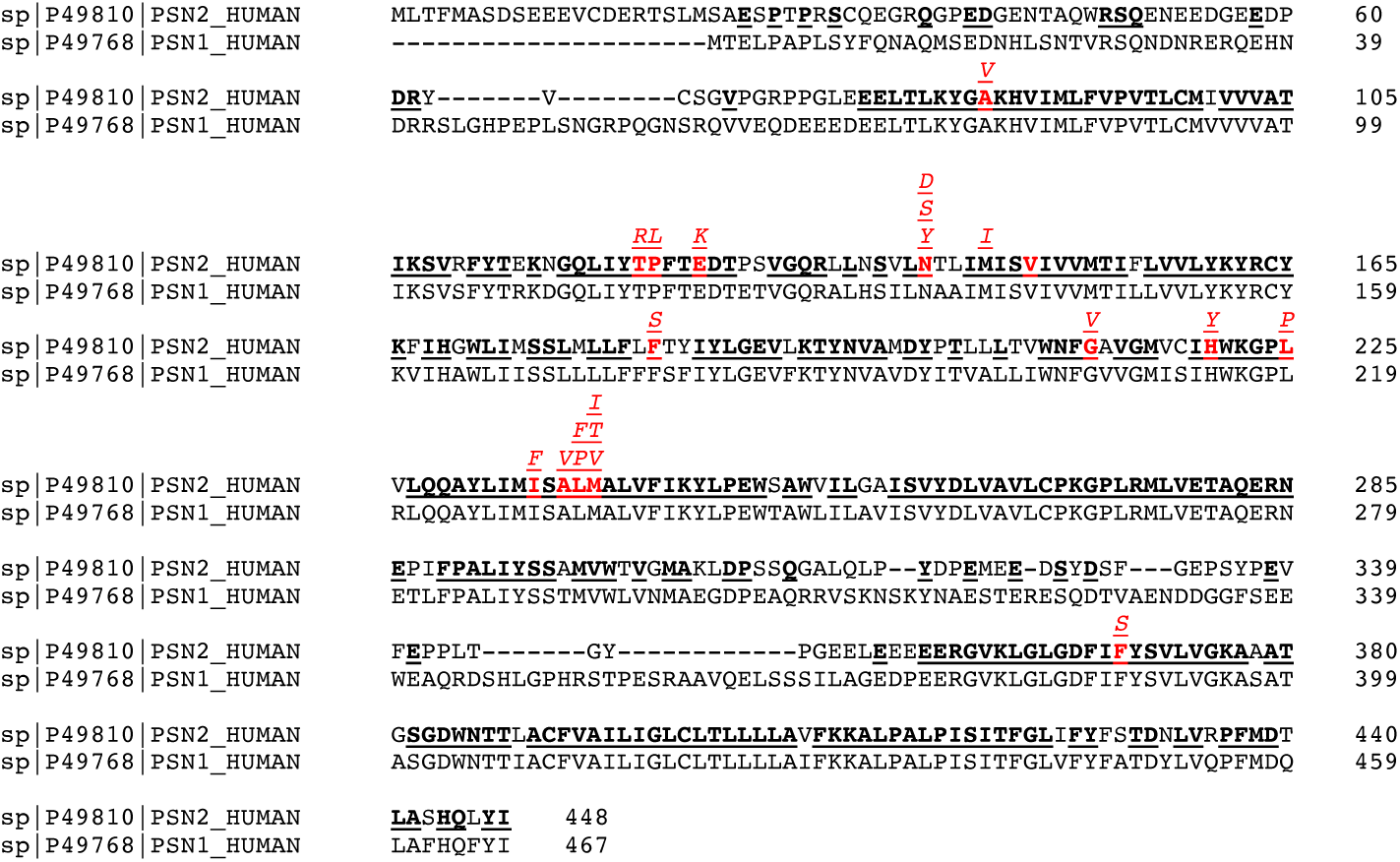
Schematic depicting the human PSN2 amino acid sequence (top row) and corresponding human PSN1 amino acid sequence (bottom). Homologous amino acid residues are bolded and underlined. Location of amino acids with known homologous variants in both PSN1 and PSN2 are bolded in red with the amino acid substitutions shown above.

**FIGURE 2.**
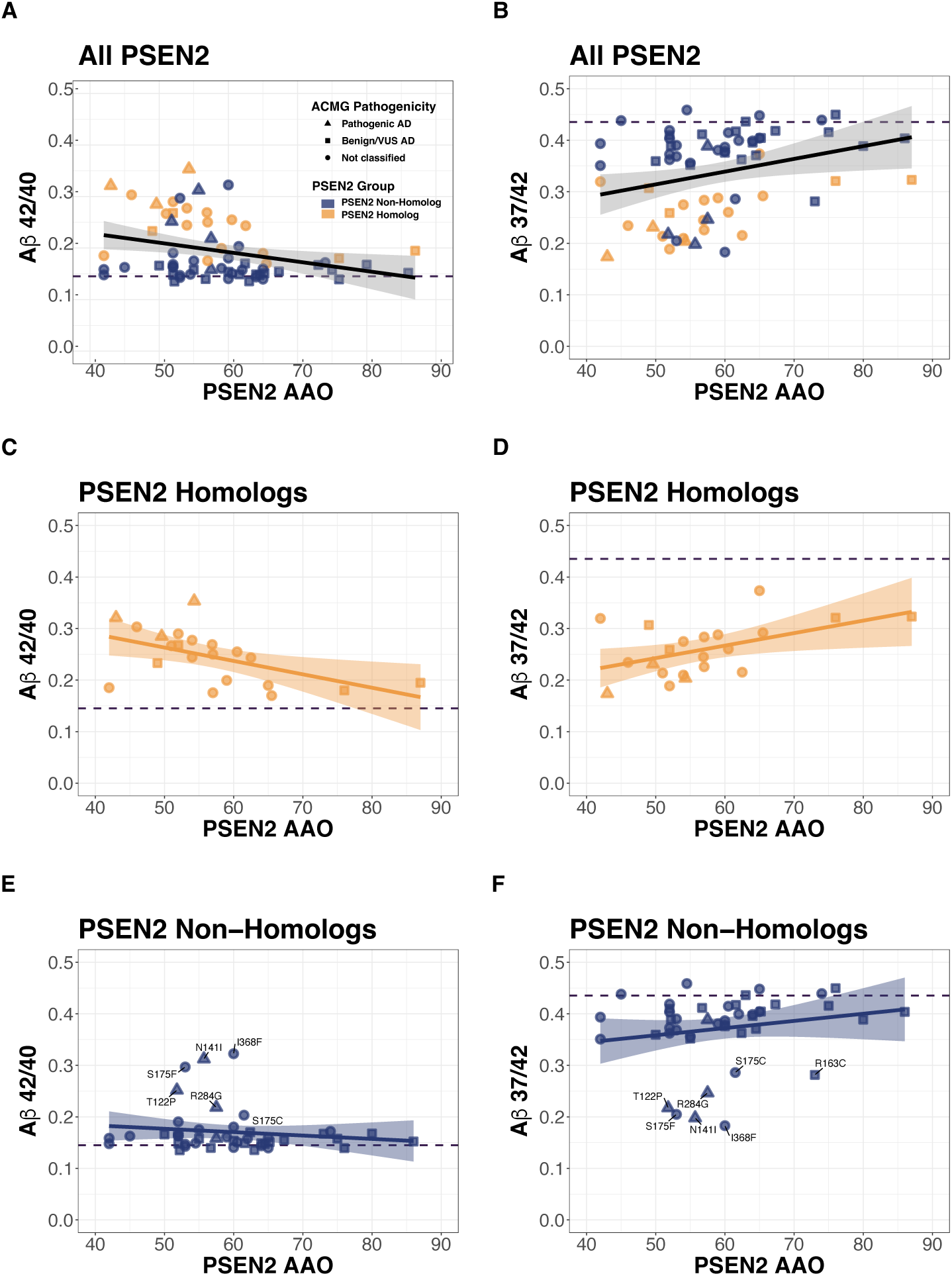
Correlation between Aβ42/40 and Aβ 37/42 cellular production ratios and AAO across all *PSEN2* variants (**A,B**), across only *PSEN2* variants with a known PSEN1 homolog (**C,D**; orange), and across only *PSEN2* variants that lack a PSEN1 homolog (Non-Homologs; **E,F**; blue). Pathogenicity classifications, presented in Table 1, for each of the *PSEN2* variants are depicted as triangles (Pathogenic AD), squares (Benign/VUS AD), or circles (Not Characterized). Wildtype *PSEN2* ratios for cell-based Aβ42/40 and Aβ37/42 are depicted as black dashed horizontal lines. The shaded area each of the linear fit lines represents the 95% confidence interval.

Notably, however, a cluster of non-homologous *PSEN2* variants demonstrated Aβ42/40 and Aβ37/42 ratios that were similar to those previously observed in *PSEN1* variants that were clearly pathogenic, suggesting that this small set of non-homologous *PSEN2* variants (T122P, S175C, S175F, N141I, and I368F) may be pathogenic despite the absence to date of a known, corresponding pathogenic mutation in *PSEN1*.

### 3.3 Comparing homologous PSEN1 and PSEN2 variants

We next compared AAO between *PSEN1* and *PSEN2* variants. We observed an average AAO of 58.4 yo, across all known *PSEN2* variant carriers, whereas *PSEN1* variant carriers bearing homologous mutations had an earlier average AAO of 40.1 yo (difference of 18.3 years; t(32) = −6.9835 and p = 6.841e-08; **Figure 3A**). There was a strong positive correlation between the AAO of *PSEN2* variants and the AAO of their *PSEN1* variant homologs (r(19) = 0.473, p = 0.030; **Figure 3B**), with a median shift of 21.6 years (range: −10, 32.5) later AAO in *PSEN2* variants compared to homologous variants in *PSEN1*. Consistent with this finding, for a given Aβ42/40 or Aβ37/42 ratio, we observed that *PSEN2* variants have a later AAO compared to homologous *PSEN1* variant (Aβ42/40: B (SE) = 0.004 (0.002) and p = 0.045; Aβ37/42: B (SE) = −0.004 (0.001) and p = 0.018; **Figure 3C,D**).

**FIGURE 3.**
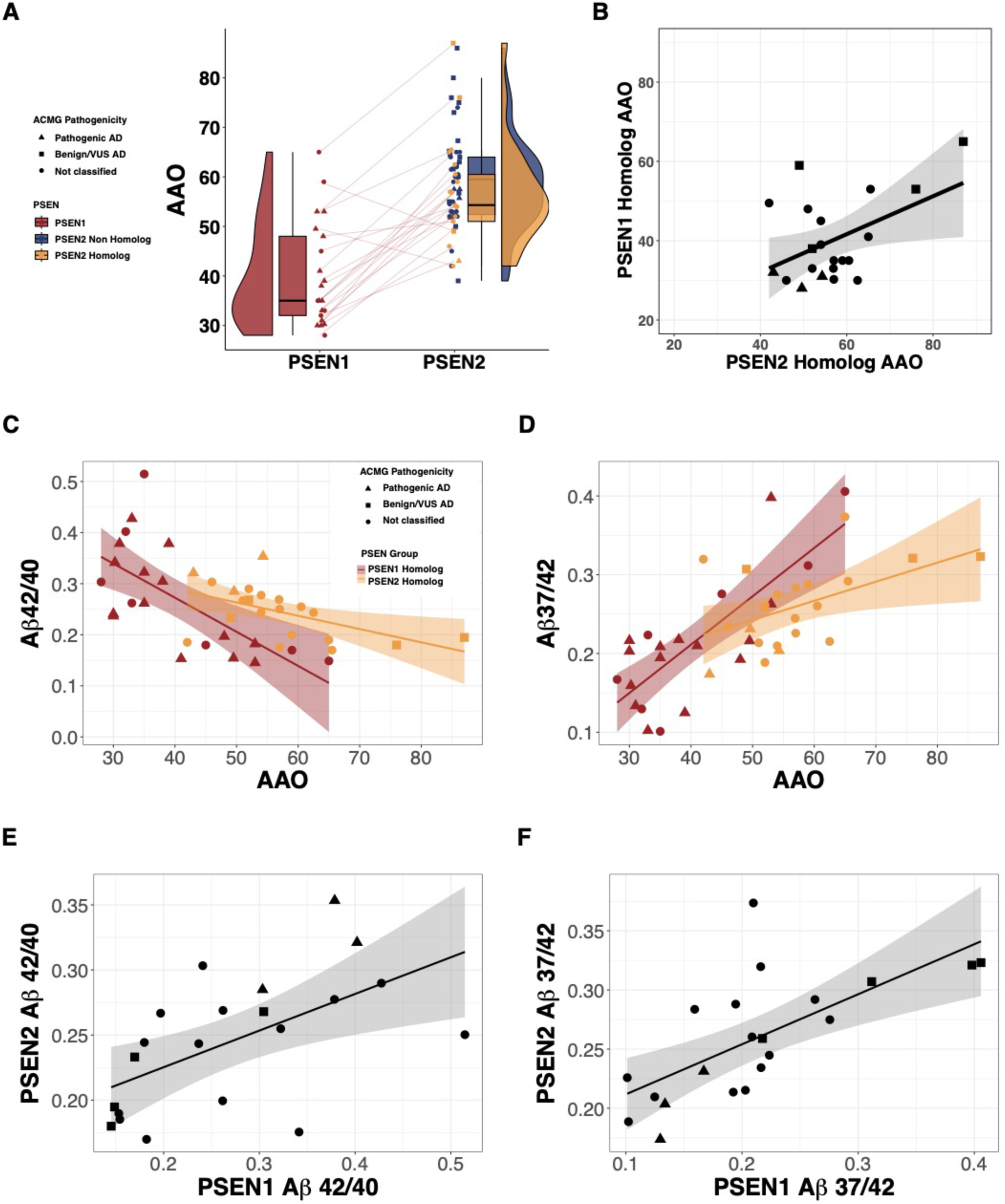
(A) Relationship between age of clinical symptom onset (AAO) in *PSEN1* (red), in *PSEN2* variants with a known *PSEN1* homolog (blue), and in *PSEN2* variants lacking a *PSEN1* homolog (orange). The boxes map to the median, 25th and 75th percentiles, and the whiskers extend to 1.5 × interquartile range (IQR). The raincloud plots illustrate kernel probability density (i.e. the width of the shaded area represents the proportion of the data located there). Lines connect a *PSEN2* Homolog variant with its corresponding homologous *PSEN1* variant and show relative shift in AAO. Pathogenicity classifications, presented in Table 1, for each of the *PSEN* variants are depicted as triangles (Pathogenic AD), squares (Benign/VUS AD), or circles (Not Classified**). (B)** Relationship between AAO of *PSEN2* variants and AAO of corresponding homologous *PSEN1* variants. **(C, D)** Correlation between cell-based Aβ42/40 (**C**) and Aβ37/42 (**D**) ratios and AAO in *PSEN2* Homolog (yellow) and *PSEN1* homologous (red) variants. **(E, F)** Relationship between AAO of *PSEN2* homologous variants and cell-based Aβ42/40 (**E**) and Aβ37/42 (**F**) ratio. The shaded area around the linear fit line represents the 95% confidence interval.

### 3.4 Relationship between cell-secreted Aβ levels in PSEN1/2 Homologs

Lastly, we compared how homologous changes in *PSEN*1 and *PSEN2* impact the production of Aβ species in our cell culture system. We observed that Aβ42/40 and Aβ37/42 production ratios were correlated between *PSEN1* variants and their *PSEN2* homologs (Aβ42/40: r = 0.580, p = 0.006 and Aβ37/42: r = 0.678, p = 0.0007; **Figure 3E,F**), suggesting that *PSEN2* variants had similar biochemical effects on γ-secretase conformation and thus proteolytic function as did the homologous mutation in *PSEN1*.

## DISCUSSION

Though it is widely recognized that numerous pathogenic variants leading to ADAD exist in *PSEN2* (*2, 16, 24*), the pathogenicity of many *PSEN2* variants remains uncertain(*38*), as does the extent to which corresponding changes between the highly similar PSN1 and PSN2 proteins may lead to similar pathogenicity, AAO, and patterns of Aβ production. Here we examined a comprehensive set of *PSEN2* pathogenic variants, including variants with uncertain pathogenicity, in order to elucidate these issues. Broadly, we observed that, similar to recent work characterizing *PSEN1* variants (*24, 27, 30, 39*), there were significant associations between the patterns of Aβ production across *PSEN2* variants and AAO. However, follow-up analyses clearly suggested that *PSEN2* variants for which there is a known, corresponding variant in *PSEN1* are more likely to have abnormal Aβ production patterns that strongly correlate with AAO. Aside from a small subset of *PSEN2* variants without a known *PSEN1* homologous variant that had Aβ42/40 ratios likely within a pathogenic range, most *PSEN2* variants lacking *PSEN1* counterparts had Aβ42/40 ratios close to those of wild-type PSN2, arguing against their pathogenicity. Together, these results support consideration of whether a *PSEN1* homolog variant is known when assessing the pathogenicity of a *PSEN2* variant of uncertain significance. Interpreted in another way, these results also suggest that there may be qualitative biochemical differences between *PSEN2* variants that have a known *PSEN1* counterpart (“homologous *PSEN2* variants”) and those that lack a known *PSEN1* counterpart (“non-homologous *PSEN2* variants”).

Intriguingly, we observed that homologous *PSEN1* and *PSEN2* variants had strongly correlated Aβ42/40 and Aβ37/42 ratios, indicating that the corresponding amino acid substitution in each presenilin may have largely similar biochemical effects on γ-secretase processivity. However, in comparing AAOs across *PSEN2* and *PSEN1* homologs, we observed a shift of 18.3 years later for *PSEN2* variants compared to their *PSEN1* counterparts. While this latter observation is consistent with what has previously been described in the literature (*24, 25*), the contrast between similar cell-based patterns of Aβ production but substantially shifted AAO between *PSEN1* and *PSEN2* homologs underscores the need for further study regarding the differences between PSN1 and PSN2 under normal physiological conditions. Several possibilities exist to explain these differences, including subtly different conformations (folding) of PS1 and PS2 in the γ-secretase complex, differential efficiency of incorporation of PSN1 vs PSN2 into the pool of active γ-secretase complexes (as previously described in Liu et. al. (*40*)), differential trafficking of PSN1 and PSN2 within the cell, and potential differences in the half-life of the PSN1 as compared to the PSN2 protein. These and other biochemical differences may explain the observation that PS2 is less proteolytically active than PS1 as an intramembrane protease and therefore less commonly the site of AD-causing mutations.

Because it relies on over-expression, the cell model system used here is not well-suited to assess the differential transfection efficiency of *PSEN1* and *PSEN2*, a limitation of the present study. Another important limitation is the relative lack of *in vivo* clinical and AD biomarker data available for *PSEN2* carriers as a group. The relative infrequency of *PSEN2* variants and the absence of a suitably large cohort of *PSEN2* variants within the Dominantly Inherited Alzheimer’s Network (DIAN) (*41*) studies makes it difficult to definitively assess the potential pathogenicity of most *PSEN2* variants with respect to AD cognitive and pathological changes. This contrasts with prior work from our group characterizing a large number of *PSEN1* pathogenic variants, where we were able to correlate a diverse set of *in vivo* clinical, cognitive, biofluid, and neuroimaging data with cell-based assessments of γ-secretase function (*27*).

Despite these limitations, the findings here suggest that the relatively simple cellular model system used here is useful for characterizing newly discovered *PSEN2* variants and making decisions about the inclusion of particular variants in ADAD clinical trials. Additionally, the characterization of a comprehensive set of *PSEN2* variants in the same cellular system is valuable for comparing AAO and pathologic changes among *PSEN2* variants. Most importantly, the present results strongly support the consideration of the existence of a *PSEN1* homologous variant when assessing the pathogenicity and estimated AAO for a known or new *PSEN2* variant.

## ACKNOWLEDGMENTS

This work was funded by NIH Grants R01AG071865 (JPC, LL, DJS) and RF1AG079569 (JPC and LL) and the Davis APP program at Brigham and Women’s Hospital (DJS, JPC, and LL). SAS was supported by an NIH grant (K01AG084816) and a post-doctoral fellowship award from the Alzheimer’s Association (US).

