## Supplemental Material for "The pathogenicity of *PSEN2* variants is tied to Aβ production and homology to *PSEN1*"

| Pathogenicity | Frequency (N) |
|---|---|
| **PSEN2 Non-Homolog (N = 53)** | |
| **AD : Not Classified[3]** | 34.0 (18) |
| **AD : Benign[2]** | 30.2 (16) |
| **AD : Uncertain Significance[2]** | 7.5 (4) |
| **AD : Likely Pathogenic[1]** | 5.7 (3) |
| **FTD : Not Classified[3]** | 5.7 (3) |
| **AD : Benign, FTD : Not Classified[3]** | 1.9 (1) |
| **AD : Likely Benign[2]** | 1.9 (1) |
| **AD : Pathogenic[1]** | 1.9 (1) |
| **AD : Uncertain Significance, FTD : Not Classified[2]** | 1.9 (1) |
| **bvFTD : Not Classified[3]** | 1.9 (1) |
| **Dementia : Not Classified[3]** | 1.9 (1) |
| **MCI : Not Classified[3]** | 1.9 (1) |
| **PD : Not Classified[3]** | 1.9 (1) |
| **svPPA : Not Classified[3]** | 1.9 (1) |
| **PSEN2 Homolog (N = 21)** | |
| **AD : Not Classified[3]** | 57.1 (12) |
| **AD : Uncertain Significance[2]** | 14.3 (3) |
| **AD : Pathogenic[1]** | 9.52 (2) |
| **AD : Benign[2]** | 4.8 (1) |
| **AD : Likely Pathogenic[1]** | 4.8 (1) |
| **AD : Not Classified, DLB : Not Classified[3]** | 4.8 (1) |
| **Atypical Dementia : Not Classified[3]** | 4.8 (1) |
| **PSEN1 Homolog (N = 21)** | |
| **AD : Pathogenic[1]** | 52.4 (11) |
| **AD : Not Classified[3]** | 33.3 (7) |
| **AD : Likely Pathogenic[1]** | 14.3 (3) |

**SUPPLEMENTAL TABLE 1.** Frequency (N) for ACMG-AMP pathogenicity classification of *PSEN2* and *PSEN1* variants. Pathogenicity classification was sub-classified into Pathogenic AD[1] (Pathogenic AD or likely pathogenic AD), Benign or VUS AD[2] (benign AD or uncertain significance AD), or Not Classified[3] (AD, Atypical Dementia, DLB, svPPA, MCI, PD, FTD, and Dementia: Not classified) groups for further group analyses (Supplemental Figure 1).

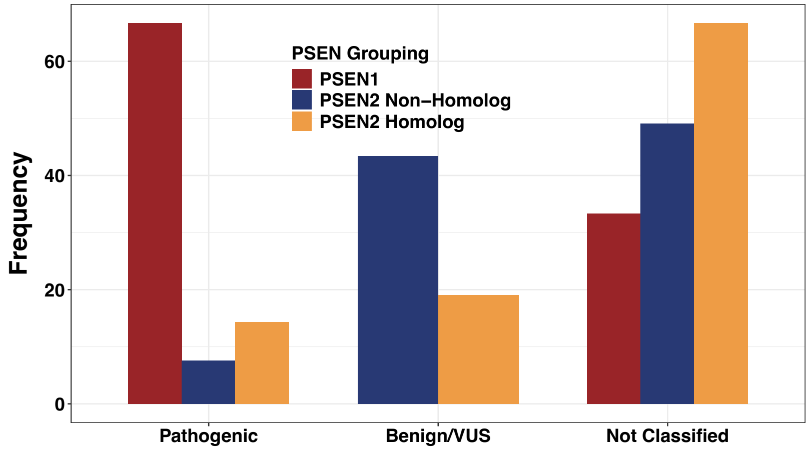


**SUPPLEMENTAL FIGURE 1.** Frequency of variants classified as Pathogenic AD (Pathogenic AD or likely pathogenic AD), Benign/VUS) AD (Benign AD or variant of unknown significance AD), or Not classified (all other classifications; see Table 1) within *PSEN1* Homolog variants (red), *PSEN2* Non-Homolog variants (blue), and *PSEN2* Homolog variants (orange).
